# Detection of circulating extracellular mRNAs by modified small RNA-sequencing analysis

**DOI:** 10.1101/507681

**Authors:** Kemal M. Akat, Youngmin A. Lee, Arlene Hurley, Pavel Morozov, Klaas E.A. Max, Miguel Brown, Kimberly Bogardus, Anuoluwapo Sopeyin, Kai Hildner, Thomas Diacovo, Markus F. Neurath, Martin Borggrefe, Thomas Tuschl

## Abstract

Extracellular mRNAs (ex-mRNAs) potentially supersede extracellular miRNAs (ex-miRNAs) and other RNA classes as biomarkers. Here, we present a comprehensive extracellular RNA (exRNA) study in human blood circulation based on conventional small RNA-sequencing (sRNA-seq) and sRNA-seq after T4 polynucleotide kinase (PNK) end-treatment of total exRNA isolated from serum and platelet-poor EDTA, ACD, and heparin plasma. Applying strict criteria for read mapping and annotation, we found that compared to conventional sRNA-seq PNK-treatment increased the detection of informative ex-mRNAs reads up to 50-fold. Based on captured ex-mRNAs from healthy individuals, we concluded that the exRNA pool is dominated by hematopoietic cells and platelets, with additional contribution from the liver. About 60% of the 15- to 42-nt long reads originated from the coding sequences, in a pattern reminiscent of ribosome-profiling studies for high abundance transcripts. Blood sample type had a considerable influence on the exRNA profile. The number of detected distinct ex-mRNA transcripts ranged from on average ~350 to 1100 in the different plasma types. In serum, additional transcripts from neutrophils and hematopoietic cells increased this number to ~2300. For EDTA and ACD, in particular, we found evidence of destabilization of mRNA and non-coding RNA ribonucleoprotein complexes. In a proof-of-concept study, we compared patients with acute coronary syndrome (ACS) to healthy controls. The improved tissue resolution of ex-mRNAs after PNK-treatment enabled us to detect a neutrophil-signature in ACS that escaped detection in an ex-miRNA analysis. Thus, ex-mRNAs provide superior resolution for the study of exRNA changes in vivo and ex vivo. They can be readily studied by sRNA-seq after T4 PNK end-treatment.

## Introduction

Extracellular RNAs (exRNAs) in biofluids were described as early as the first half of the 20th century^1^ but underwent a more recent renaissance with the detection of circulating miRNAs^2^. Despite the high nuclease activity in biofluids, miRNAs (ex-miRNAs) remain detectable due to protection by tightly bound RNA-binding proteins and/or inclusion in microvesicles^2–5^. In recent years, especially with the advancement of RNA-sequencing (RNA-seq), an extensive body of research accumulated regarding the role of extracellular miRNAs in a broad range of medical conditions and cardiovascular diseases, including advanced heart failure^6^ and myocardial infarction^7^.

Ex-miRNAs are remarkably stable in circulation, and we recently showed that distinct ex-miRNA signatures can be followed for months^8^. However, a general limitation of ex-miRNAs is the relatively low number of miRNA genes with only few tissue specific members^9^. Alterations of ex-miRNAs are often difficult to interpret biologically as they either affect ubiquitously expressed or low-abundance miRNAs without a clearly identifiable source tissue. In contrast, the number of mRNA genes in the human genome is at least an order of magnitude higher^10^ providing a much better tissue and functional resolution for physiological conditions or disease states. While RNA-sequencing (RNA-seq) potentially offers the most comprehensive interrogation of ex-mRNAs and their changes lack of robust protocols and challenges in the analysis of fragmented, short reads hampered their study.

Technical challenges in exRNA profiling encompass the very low amounts of RNA in body fluids, and the influence of anticoagulants used for blood collection, increasing the likelihood for batch effects or spurious findings^11,12^. The type of blood sample used for RNA isolation can substantially influence the stability of certain ribonucleoprotein (RNPs) complexes and associated RNAs. A striking example for differential stability of RNPs with different anticoagulants is the loss of 5’ tRNA fragments using magnesium-ion-chelating EDTA or citrate salts for blood collection^6,13^. While it seems likely that these routinely used chelators for blood collections will impact the stability of other extracellular RNPs, the overall extent in which the sample types influence the exRNA profile remains unknown.

By design sRNA-seq cDNA protocols enrich for miRNAs, which carry 5 ‘ phosphate and 3 ‘ hydroxyl groups. However, in body fluids other classes of RNAs, including potentially mRNAs, most likely exist as degradation products due to the high nuclease activity^8^. RNA degradation products possess 5’ OH ends as well as 2’ or 3’ phosphate or 2’,3’ cyclic phosphate termini. These termini are incompatible with sRNA-seq, and fragments of those RNAs will largely escape detection. Enzymatic treatment of RNA ends by T4 polynucleotide kinase (PNK) rescues RNA fragments devoid of the necessary termini and has been used for different RNA-seq based applications including exRNA studies^14,15^. However, an effect on ex-mRNA capture has not been shown thus far.

Here, we used a recently published RNA isolation protocol that quantitatively recovers exRNAs^8^, and combine T4 PNK RNA end-modification with sRNA-seq and stringent read annotation criteria to demonstrate effective and informative capture of ex-mRNAs. We investigated blood samples with different commonly used anticoagulants to identify confounding factors, and finally tested the potential of ex-mRNAs in a proof-of-concept cohort of patients presenting with an acute coronary syndrome.

## Methods

### Sample procurement

Blood was collected from healthy volunteers and from patients evaluated for acute coronary syndrome at The Rockefeller University and Mannheim University Medical Centre, respectively, by the first author. Human tissue samples for bulk mRNA-seq were obtained from the National Disease Research Interchange (Philadelphia), or from biopsies or discarded surgical waste. Sample procurement was approved by the institutional review boards of all participating institutions. All participants gave written informed consent, and the studies were approved by the IRBs of the participating institutions.

### RNA isolation

ExRNA was isolated from 425 μl cell-free serum or platelet-depleted plasma using a customized RNA isolation protocol developed to minimize residual nuclease activity^8^; the RNA was purified using silica columns. Cellular or tissue total RNA was extracted using TRIzol with an additional phenol/chloroform extraction step and concentrated by alcohol precipitation.

### PNK treatment of total exRNA

After elution from the silica column, half of the isolated total exRNA was used directly for sRNA-seq, and the other half treated with T4 PNK in a total reaction volume of 20 μl for 30 min at 37 °C followed by re-purification and elution of the PNK treated RNA using the same silica column, and then subjected to sRNA-seq library preparation.

### Small RNA-seq

sRNA-seq cDNA library preparation was done as described^16^ but size selecting from 19- to 45-nt. Long mRNAseq of cells and tissues was done using the Illumina Stranded mRNA-seq TruSeq protocol following the manufacturer’s instructions. Sequencing was conducted in the Genomics Core Facility at The Rockefeller University.

### Bioinformatics analysis

#### Read annotation

Read processing and annotation for small RNA-seq of serum and plasma samples was done as described^17^ with modifications for PNK-treated samples. Long RNA-seq reads from tissues or cells were aligned to the human genome build 38 using the STAR aligner^18^ and quantified using the featureCounts^19^ program based on Ensembl release 82.

#### Data analysis and statistics

Differential analysis, clustering, and other downstream analyses were done in the R statistical language and Bioconductor packages. Other statistical tests are indicated in text and figures where appropriate. If not stated otherwise, results with a p value < 5% were considered significant (Benjamini-Hochberg adjusted for all RNA-seq comparisons).

#### Tissue specificity score

An RNA-seq expression atlas comprised of representative tissues was used to calculate a tissue-specificity score to identify the source tissue of circulating mRNAs^10^.

### Clinical laboratory parameters

Standard clinical laboratory assays were performed by the Central Laboratories of the University Medical Centre Mannheim, Mannheim, Germany, and Memorial Sloan Kettering Cancer Centre, New York, NY, USA.

## Results

### PNK treatment of exRNA improves the capture of mRNA fragments by small RNA-seq

To test if PNK treatment improves capture of ex-mRNA fragments we performed sRNA-seq comparing untreated to PNK-treated total exRNA input from the same donors. Different anticoagulants were used to assess their influence on the exRNA profile (Fig. 1).

**Fig. 1.**
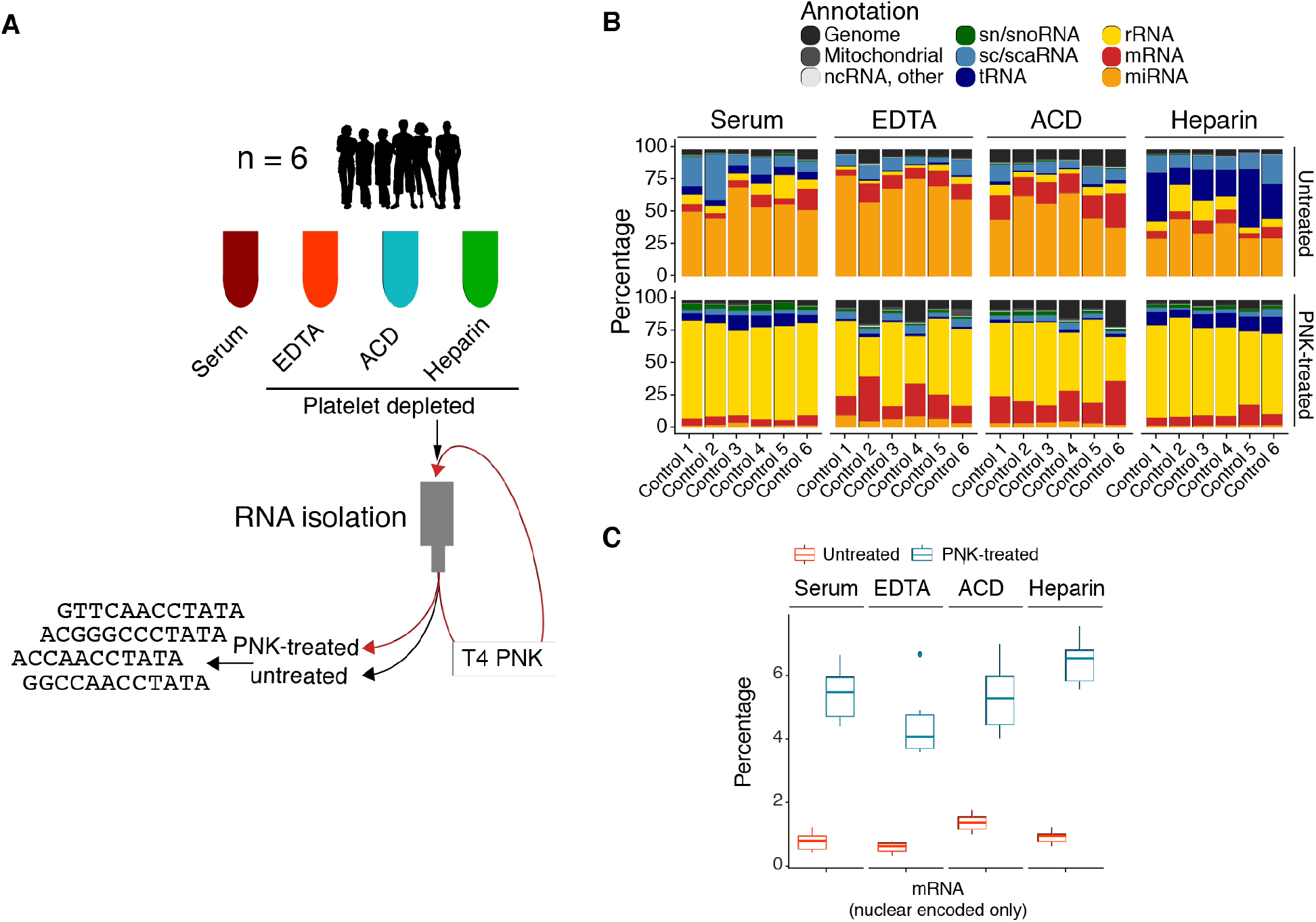
Treatment of total extracellular RNA with T4 polynucleotide kinase (T4 PNK) followed by small RNA-sequencing (sRNA-seq). (A) Total RNA was isolated from 450 μl serum or platelet-depleted EDTA, acid citrate dextrose (ACD), and heparin plasma from 6 healthy individuals and purified using silica-based spin columns. Half of the RNA was treated with T4 PNK and re-purified (PNK-treated) and multiplexed sRNA-seq libraries were prepared separately for the untreated (libraries 1 and 2) and end-treated RNA (libraries 3 and 4). (B) Differences in read annotation in the four sample types for untreated RNA and PNK-treated RNA using initial annotation settings (up to 2 mismatches, multi-mapping). (C) Differences in nuclear mRNA capture between untreated and PNK-treated RNA using final annotation criteria (no mismatch and up to two mapping locations).

Plasma and serum were collected from six healthy volunteers. Collection tubes for plasma samples contained the divalent-metal-ion-chelating EDTA and ACD salts or the polyanion heparin. All plasma samples were platelet-depleted, and total exRNA was recovered by our recently published isolation protocol which preserves RNA integrity and quantitatively recovers exRNA^8^. Multiplexed sRNA-seq libraries were generated minimizing batch effects (libraries 1-4; Fig. 1, Supplementary Data 1).

More than 95% of the processed reads were 12- to 42-nt in length. Such short reads impose challenges for confident transcript assignment due to multi-mapping. For conventional sRNA-seq, i.e. miRNA studies, this is minimized by hierarchical mapping and requiring a minimum read length of 16-nt^17^. Hierarchical mapping ensures that more abundant RNAs like rRNAs and tRNAs take precedence over less abundant classes like mRNAs and miRNAs if a sequence matches to more than one RNA class. To arrive at a comprehensive assessment, we initially retained reads <16-nt. With that, over 80% of reads mapped to established classes of human RNAs and human genome with the expected enrichment for miRNAs in the untreated samples (Fig. 1B). The most apparent difference after PNK-treatment was the increase in the rRNA fraction. A residual 3-15% of reads mapped to the *E. coli* genome, and ~1% to bacterial expression plasmids and diatoms (Supplementary Data 1). Bacterial RNA is a common contaminant in recombinantly produced enzymes used for library preparation and residual diatom RNA exists in commercial silica matrices used for nucleic acid isolation. In standard RNA-seq applications using higher amounts of input RNA, these sequences do not influence the results but they can contribute a sizeable fraction of sequence reads in low input samples like body fluids^6,8^.

Ex-mRNA reads comprised 6.5 to 20% with some enrichment after PNK-treatment in EDTA and ACD plasma but not in heparin plasma or serum (Fig. 1B). Further review of read alignments, however, showed that untreated samples collected more mRNA reads with 1 or 2 mismatches, i.e. inflating the mRNA read count by inclusion of low-confidence reads (Supplementary Data 2). As expected, reads <15-nt had a high fraction of multi-mapping (Supplementary Fig. 1). Therefore, our final ex-mRNA analysis was restricted to perfectly mapping reads (0 mismatch) 15-nt or longer with at most two mapping locations. The latter was necessary to account for the identical coding sequences of the hemoglobin paralogs HBA1 and HBA2 that would otherwise be underrepresented.

Using these annotation criteria PNK-treatment unambiguously increased the percentage of ex-mRNA reads and even more the number of unique transcripts captured. Compared to untreated samples, in PNK-treated samples the mRNA read count increased ~4-fold in ACD samples and ~9-fold in all other sample types (Fig. 1C). Requiring 5 unique reads per mRNA and donor sample, we captured an average (min, max) of 2313 (452, 4634), 583 (162, 1192), 350 (75, 625), and 1108 (591, 1760) distinct mRNA transcripts in serum, EDTA plasma, ACD plasma, and heparin plasma samples, respectively. This compared to only 46 (2, 182), 33 (1, 86), 27 (5, 70), and 43 (0, 140) distinct mRNAs in the corresponding untreated samples (P value < 8e-09, Wilcoxon rank sum test), representing a 13- to 50-fold increase.

### Ex-mRNAs in circulation originate mostly from the coding sequences and not UTRs

It has been previously reported, that ex-mRNAs in cell culture media mostly originate from the 3 ‘ UTR of mRNA transcripts^20^. Review of read alignments in our study, however, indicated that most of the ex-mRNA reads originated from the transcript coding sequence (CDS), a pattern that was only observable in PNK-treated samples due to better transcript coverage. Read distribution and read length were reminiscent of ribosome-profiling data, which indicated that ex-mRNA fragments are ribosome protected and circulate as polysome or monosome complexes. This observation was confirmed by a metagene analysis that was based on an average of 12,789 to 16,486 ex-mRNA transcripts depending on sample type. This showed that ~60% of the reads originated from the CDS and ~30% from the 3’ UTR (Fig. 2).

**Fig. 2.**
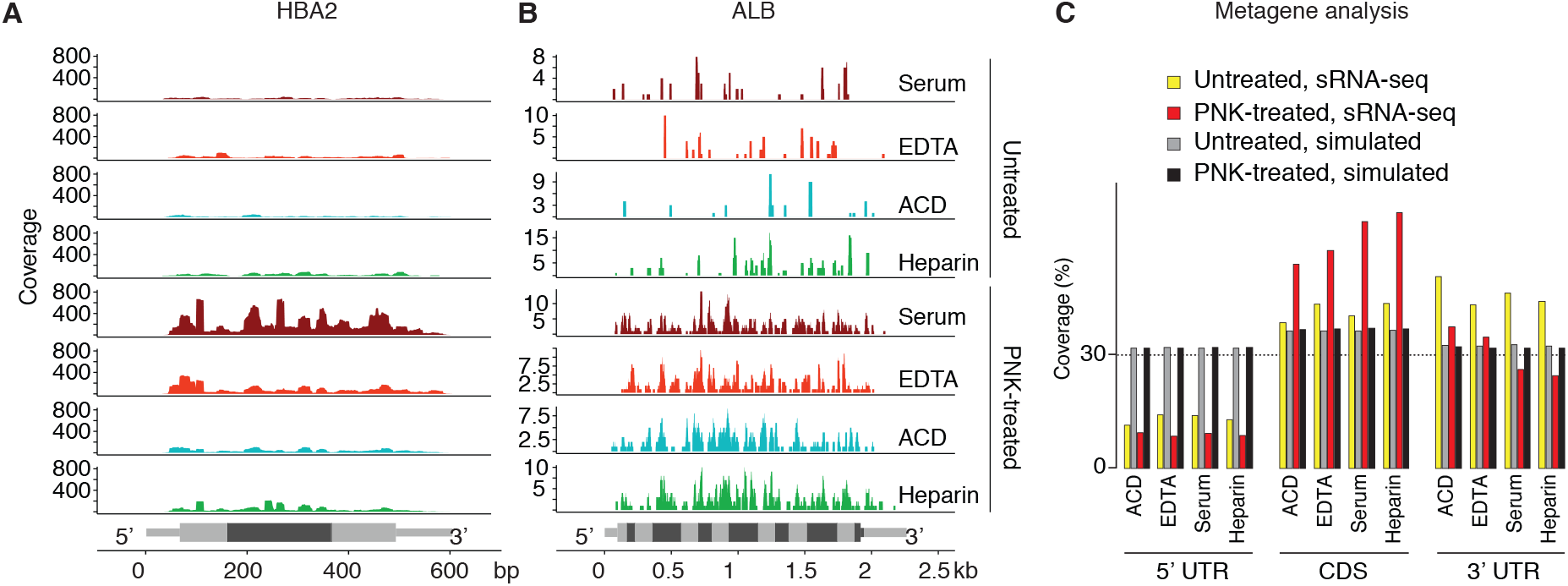
Read distribution of reads across mRNA transcripts. (A, B) Read coverage for the hemoglobin A2 transcript (A) and the albumin transcript (B) by sample type for untreated (upper rows) and T4 PNK end-treated (lower rows) samples. Exon boundaries (HBA2: 3 exons, ALB: 15 exons) are indicated by alternating intensities of grey, and UTRs are distinguished from CDS by thinner bars. (C) Metagene analysis with relative read coverage (percentage) across 5’ UTRs, CDS, 3’ UTRs for untreated and PNK-treated samples as well as corresponding data obtained after 100 random simulations (across an average of 759 to 3,500 captured transcripts for untreated samples and an average of 2,750 to 16,487 captured transcripts for PNK-treated samples depending on sample type).

### Anticoagulants have a widespread effect on the exRNA profile

The anticoagulants we studied are the predominant ones used to collect blood samples in clinical practice and for research purposes. All of them influence blood cells ex vivo^21–23^, and heparin may not be removed sufficiently by common extraction protocols and as a result interfere with downstream applications^24^. This is especially relevant if patient populations are studied that often receive high doses of heparin.

We therefore next looked at how sample type influenced the measured exRNA composition for both the untreated and PNK-treated samples. We noted the previously reported destabilization of 5’ tRNA fragments in EDTA and ACD samples (Supplementary Fig. 2)^6,13^, and alterations in miRNA composition between serum and platelet-depleted EDTA plasma^12^. In an ANOVA-like comparison we observed abundance differences for 86 miRNAs in the untreated samples and of 1,458 mRNA transcripts in the PNK-treated samples between the three plasma types and serum (Supplementary Data 3 and 4). Serum generally had a higher abundance of ex-miRNAs (e.g. miR-223 and -142) and ex-mRNAs (e.g. S100A8) enriched in myeloid cells and platelets. In a gene set analysis ex-mRNAs abundant in serum were associated with inflammation and leukocyte activation whereas plasma ex-mRNAs were more related to general cellular processes like translation (Supplementary Data 5). Although there was a high degree of similarity between the exRNA profiles of EDTA and ACD plasma, as expected from their mechanism of action (Supplementary Fig. 3B), there were distinctive differences as well. For instance, EDTA plasma had increased levels of erythropoietic transcripts, i.e. miR-451(1) and hemoglobin mRNAs, compared to all other samples. ACD had 3- to 4-fold higher levels of miR-150(1), a lymphocyte-restricted miRNA, than the other sample types (Supplementary Fig. 3A, Supplementary Data 3 and 4).

The destabilizing effect of the chelating reagents, ACD and EDTA, on ribonucleoprotein complexes was not restricted to tRNAs. Both altered read coverage signatures of other RNAs. Human small nuclear RNAs U1 and U2 snRNAs are ~164-nt and ~190-nt non-coding RNAs, respectively, which assemble with proteins into small nuclear ribonucleoproteins (snRNPs). Biochemical studies demonstrated that U1 and U2 possess core structures that are relatively resistant to nuclease digestion^25^. In high magnesium conditions several U1 domains are protected from nuclease digestion whereas in low magnesium conditions, i.e. after the addition of EDTA or similar chelating reagents, only the core region remained relatively resistant to digestion. Our sRNA-seq data agreed well with these earlier observations (Supplementary Fig. 3C). In addition, the coverage of the more protected core region was 4- to 8-fold lower in EDTA and ACD plasma, respectively, than in the other two sample types. There was no difference in read coverage patterns for snRNAs U2, U4, U5, and U6 or the large ribosomal subunits, 18S and 28S, between the different sample types.

### Hematopoietic cells, platelets, and liver are the major sources of exRNAs in healthy individuals

We next sought to identify contributing tissue sources to the exRNA pool in the physiological state. We generated a polyA mRNA-seq tissue atlas comprising major human cell and tissue types and calculated a tissue specificity score (TSS)^10^ for all of the 19,810 mRNAs as defined in Ensembl release 82 (Supplementary Data 6). Genes restricted to a few tissues or cell types had a TSS greater than 3, e.g. aldolase B (ALDOB) expressed in liver and kidney, while classic marker genes like albumin (ALB) or cardiac troponin T (TNNT2) had a TSS greater than 4.

A total of 3,167 ex-mRNAs entered comparative analysis, and of those 144 had a TSS > 3 (102 >3 but <4, 42 >4; Supplementary Data 7), therefore being most informative regarding tissue of origin. About 30% of the 144 mRNAs were most abundant in neutrophils, 10% in liver, and 5% each in red blood cells (RBCs), platelets, and skeletal muscle. Conversely, when we compared the 1,000 highest expressed mRNAs for each tissue in the atlas to the 3,167 ex-mRNAs, we found a much higher fraction of the top 1,000 transcripts from RBCs, platelets, neutrophils, PBMC, and monocytes captured in circulation than from any of the other tissue (Fig. 3 and 4, Supplementary Fig. 4, Supplementary Data 8). Our annotation criteria led to the detection of certain highly tissue-specific genes from other tissues, e.g. MYBPC3 (myocardium), SFTPB (lung), or MIOX (kidney; Supplementary Fig. 4) in some serum or plasma sample types. However, the underlying reads were repetitive and short and therefore highly suggestive of annotation artefacts.

**Fig. 3.**
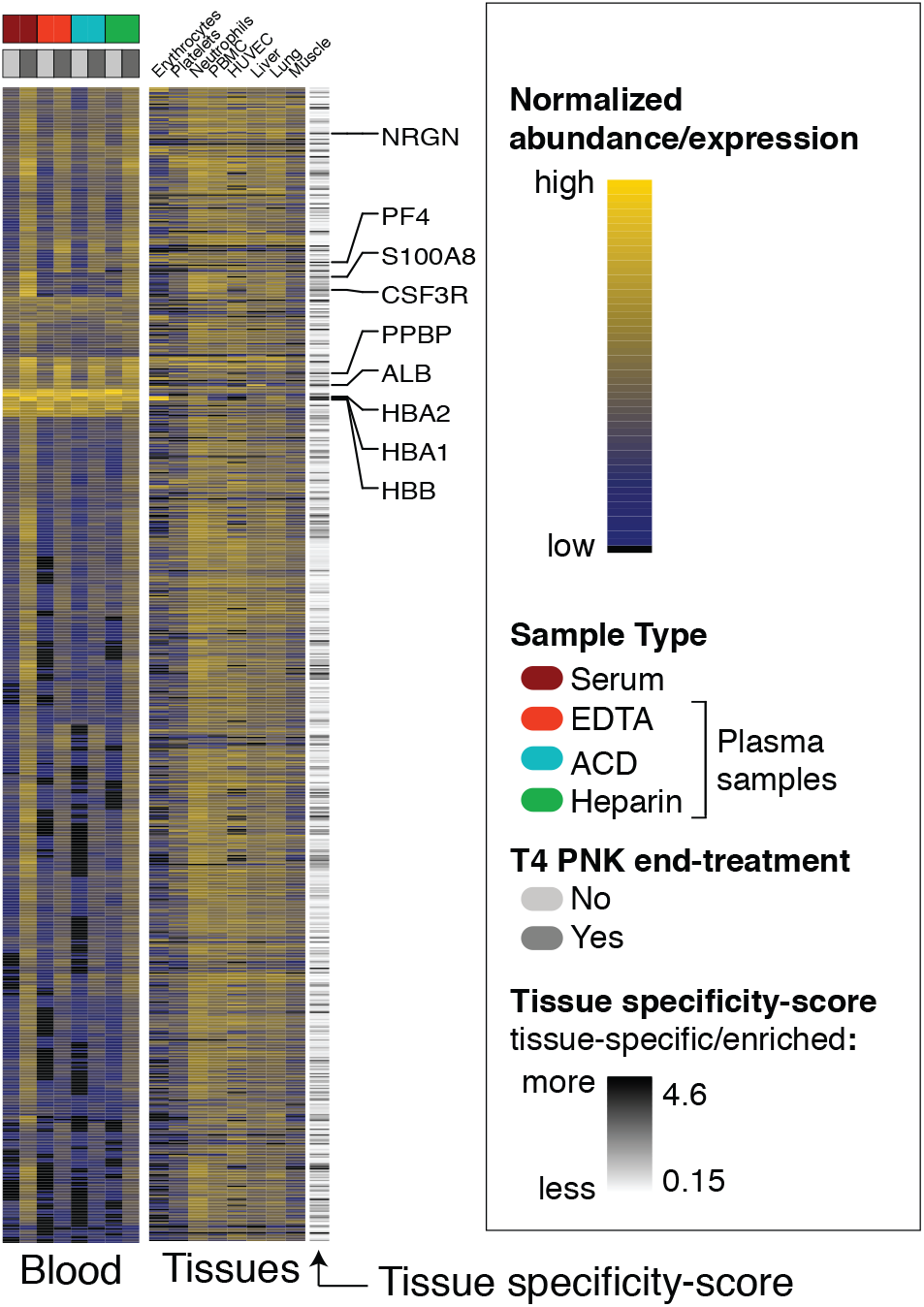
Tissue sources of circulating mRNAs (A). Heat map with the top the 821 most abundant ex-mRNAs in circulation for untreated and PNK treated (left) together with selected cells or tissues (right). Selected, tissue-specific/enriched miRNAs and mRNAs are labelled together with the tissue-specificity score.

**Fig. 4.**
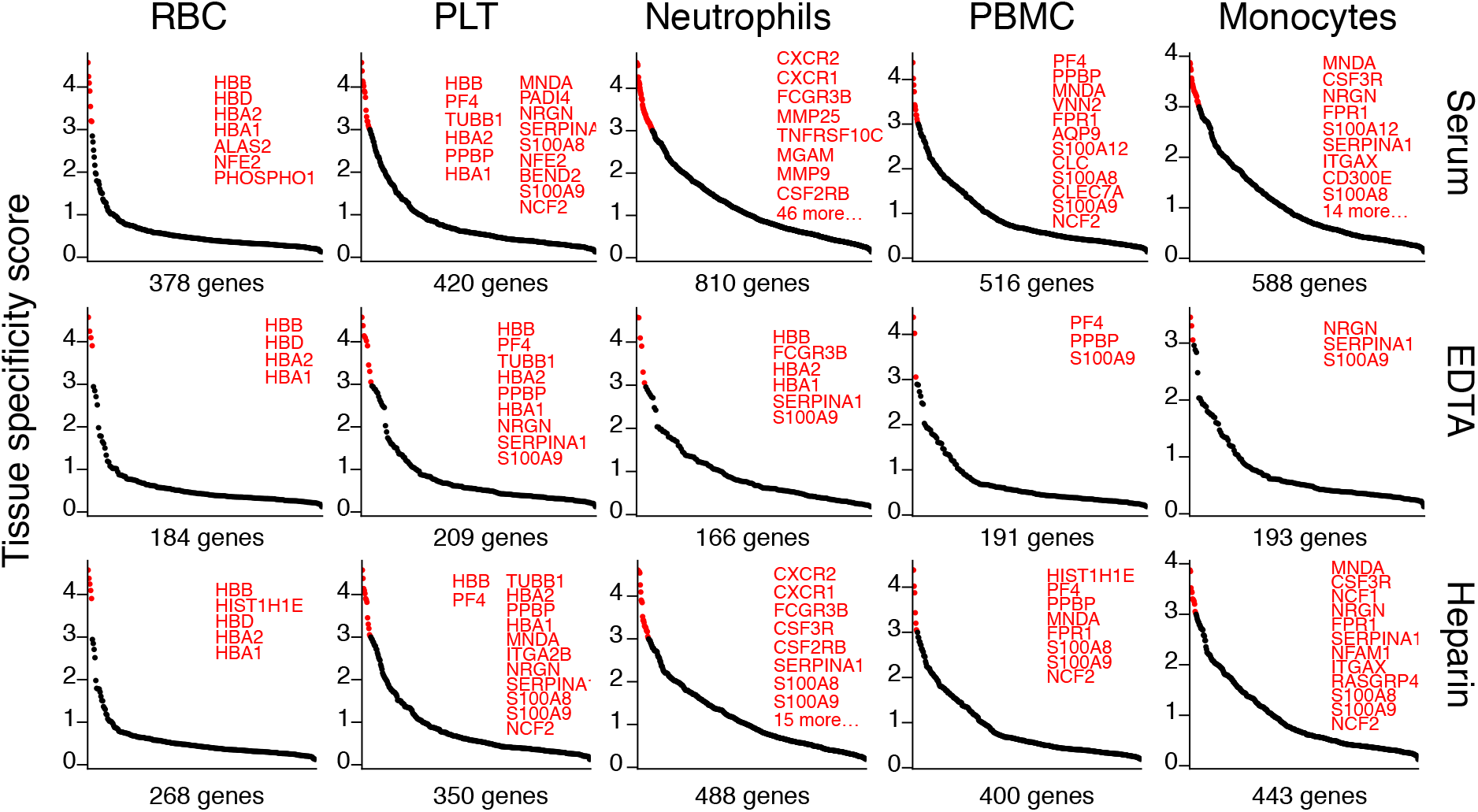
Top expressed transcripts from hematopoietic tissues captured in circulation. The 1,000 most abundant nuclear mRNA transcripts from the selected cell types that collected 5 unique or 10 total reads in at least 3 of the 6 donors per sample type were considered captured. The captured transcripts (x axis) were ordered in descending order by the tissue specificity score (TSS, y axis). Transcripts with a TSS greater than 3 were highlighted in red and listed, space permitting.

We noted, again, a clear difference between sample types. In EDTA and ACD plasma we detected 12% to 21% of the top 1,000 hematopoietic mRNAs. This percentage increased to 27% to 49% in heparin, and 38% to 81% in serum. Particularly striking was this difference for neutrophils, for which we detected 17%, 49%, and 81% of the 1,000 most highly expressed transcripts in ACD, EDTA, and serum, respectively, as ex-mRNAs (Fig. 4, Supplementary Fig. 4 Supplementary Data 8). The increase of ex-mRNAs in serum compared to the other samples is likely related to in vitro neutrophil degranulation and apoptosis during coagulation. On the mRNA level this is much more pronounced for neutrophil than platelets transcripts, of which we detected 35% in heparin and 42% in serum. Although miRNAs have been reported as markers for platelet activation^12^, our data suggest that neutrophils also contribute to coagulation-dependent miRNA abundance changes.

In summary, these results indicated that hematopoietic cells, platelets, and the liver are main contributors to the ex-mRNA profile and based on our data there was little support that other solid tissues contribute substantially.

### RNA end-modification increases the diagnostic potential of exRNA in disease

To evaluate the clinical potential of ex-mRNAs in patients we studied exRNA changes in a pilot cohort of patients with an acute coronary syndrome (ACS; n = 6) and age- and gender-matched healthy controls (n = 10; Supplementary Data 1 and 9). All patients had evidence of myocardial necrosis based on elevated cardiac troponin I levels, highly-sensitive and routinely used marker for myocardial damage. Patients with myocardial injury provided a good proof-of-concept cohort as the myocardium is one of the few tissues expressing tissue-specific miRNAs (myomirs miR-208a, -208b, and -499), which have been shown to be elevated in the circulation of these patients. In comparison to the controls the ACS group had higher white blood cell counts (Supplementary Data 9 and 10).

Because the ACS group received high doses of heparin before sample collection as part of, all patient and control samples were collected in heparin plasma to avoid any biases associated with different anticoagulants as described before. Two small RNA-seq libraries were generated from untreated (library 5) and PNK-treated (library 6) total RNA (Supplementary Fig. 5). Unsupervised hierarchical clustering of the 3’-adapter spike-in small RNAs did not separate the two groups, arguing against any potential bias due to residual heparin in the samples (Supplementary Fig. 6).

In the differential analysis 18 miRNAs were altered in the untreated samples, 11 higher and 7 lower in ACS than controls (Fig. 5A; Supplementary Data 11). The myocardium-specific miRNA miR-208b(1) in the ACS group was 17-fold higher than in the controls, the other two myocardium-specific miRNAs miR-208a (FDR 0.07%) and miR-499 (FDR 0.15%) were elevated 8-fold in ACS. These changes were in line with release due to myocardial injury and in magnitude similar to what we reported for patients in advanced heart failure^6^, and again supporting that any heparin-associated bias did not substantially influence this comparison. Individual myeloid-enriched miRNAs were elevated in ACS as well, e.g. miR-223(1), while platelet miRNAs in general were not changed (Fig. 5A).

**Fig. 5.**
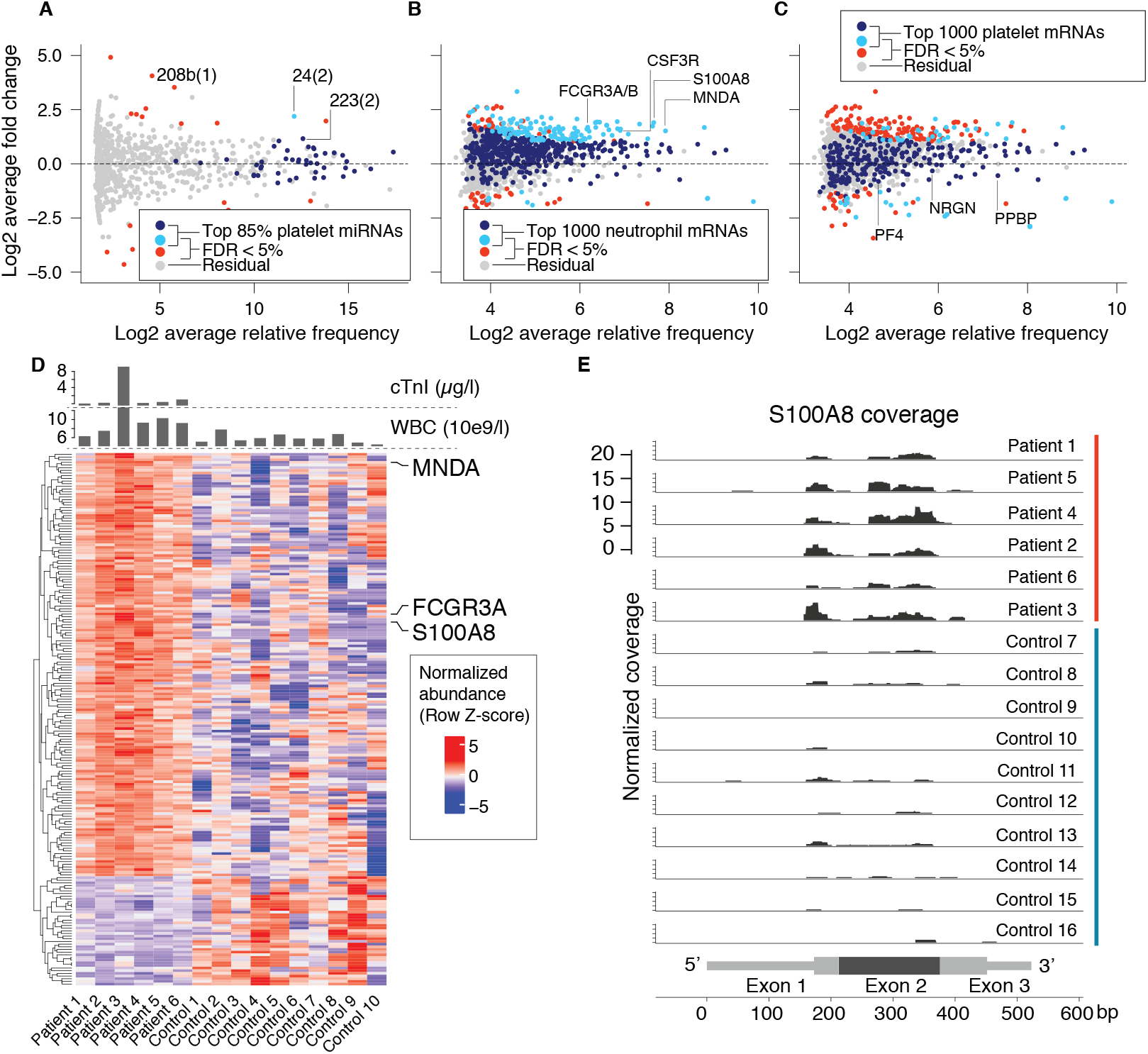
Changes in extracellular mRNAs and miRNAs in patients with ACS compared to controls. (A) MA plot of ex-miRNAs changes color coding highly expressed miRNAs in platelets defined as the top 85% miRNAs. (B, C) MA plot of ex-mRNA changes color coding highly expressed neutrophil genes (B) or platelet genes (C). Navy blue: highly expressed and FDR > 5%; orange: highly expressed and FDR < 5%; red: not highly expressed and FDR < 5%; grey: all other. Selected transcripts are highlighted: (A) myocardium specific miRNAs: 208a(1), 208b(1), 499(1); miRNAs abundant in neutrophils: 185(1), 24(2), 223(2), or miRNAs specific, 122(1), or typic, 192(1), for liver; (B) mRNA transcripts highly enriched in neutrophils or platelets (C). (D) Heat map showing altered ex-mRNAs in the ACS group compared to healthy controls. Selected mRNAs are indicated on the right. (E) RNA-seq read coverage of the 523-nt S100A8 transcript in ACS group and healthy controls (down-sampled to 600,000 reads). Transcript structure indicated at the bottom with the three exons in alternating intensities of grey, and the 5’/3’ UTRs as thin bars.

In agreement with our previous results after PNK-treatment, it improved the detection of distinct ex-mRNAs 30-fold, with an average (min, max) of 1124 (47, 4825) ex-mRNAs captured in the PNK-treated samples compared to an average of 38 (6, 313) in the untreated samples. Differential analysis identified 209 changed mRNA transcripts, 167 higher and 42 lower in ACS than controls. Most prominent was a marked increase in neutrophil transcripts in ACS (Fig. 5B, Supplementary Data 12) while platelet transcripts like the highly specific PF4 and PPBP were unchanged between the two groups (Fig. 5C). The top 6 elevated mRNA fragments in the ACS group by FDR (Fig. 5D) were IFITM2 (4.2-fold, TSS 2.25), MGAM (10-fold, TSS 4.3), CXCR2 (4.5-fold, TSS 4.1), H3F3A (3.6-fold, TSS 0.74), GCA (3.8-fold, TSS 3.2), and S100A8 (3.7-fold, TSS 3.2) all of which highly expressed in neutrophils (Supplementary Data 6) and many specifically expressed in this cell type. The reads of the released neutrophil transcripts originated again mainly from the CDS of the transcripts (Fig. 5E). In contrast to our observations with myocardium-specific miRNAs, we did not detect any myocardial mRNAs in circulation.

Taken together, these data support that ex-mRNAs a neutrophil signature in the ACS group with a release of ribosome-associated transcripts, a change not detectable on the miRNA level.

## Discussion

Here, we showed that mRNA fragments in circulation (ex-mRNAs) can be efficiently captured by T4 polynucleotide kinase (PNK) end-treatment of total extracellular RNA (exRNA) followed by sRNA-seq. Ex-mRNAs provide superior tissue and functional resolution for most conditions compared to other RNA classes because of the higher number of comparatively well annotated, highly expressed tissue-restricted transcripts. Tissue-specific ex-miRNAs, in selected cases, offer complementary information.

Ex-miRNAs have been widely studied as biomarkers in many types of diseases and conditions^6,7,26,27^. They perform well in the detection of tissue damage of organs with tissue-specific miRNAs like the liver (miR-122)^28^ or the heart (myomirs)^6,7^. Individual miRNAs alone or in combination are also used for risk prediction for chronic conditions^27^, and characteristic ex-miRNA changes have been shown to be stable over months even in the absence of detectable illness^8^. But the precise tissue-source or etiology of such differences based on the ex-miRNA profile remain unclear. Many tissues do not possess specifically-expressed miRNAs, and measurements of ubiquitously or weakly expressed miRNAs in biofluids are prone to misinterpretation.

Patients with acute coronary syndrome (ACS) represented a good benchmark population to evaluate our analysis of ex-mRNAs given the consistently reported elevations of myocardium-specific miRNAs (myomirs) in circulation^6,7^. As expected, myomirs were elevated in ACS but aside from these changes few alterations were detectable between ACS and healthy controls on the ex-miRNA level. However, the ACS group had a characteristic neutrophil ex-mRNA signature in circulation, i.e. elevated levels of neutrophil-enriched and -specific genes. Although this finding needs validation in larger cohorts and could have been confounded by the higher leukocyte count in the ACS group, the results are in line with the increasing recognition of inflammation and neutrophil activation for atherosclerotic disease. Endothelial damage and neutrophil activation have been linked to thrombus formation in animal studies^29^, and neutrophils in atherosclerotic plaques are detectable in animal models as well as human samples^30^. Irrespective of the reason for the neutrophil signature in the ACS cohort, i.e. an inflammatory response to ACS or due to higher neutrophil counts, the results clearly emphasize the superior tissue resolution of ex-mRNAs compared to ex-miRNAs. The lack of detectable myocardial ex-mRNAs in any of the samples used in this study is most likely due to the low sequencing depth of ex-mRNAs caused mainly by large of rRNA fractions but differential stability of ex-miRNAs and ex-mRNA fragments likely contributes^5^.

While our study did not address different mechanisms of exRNA release or the different compartments of exRNAs currently discussed, a few findings suggest that ex-mRNAs and probably a large part of all exRNA circulate within polysome complexes. First, in ex-mRNAs transcripts sequenced with good coverage, i.e. high abundance transcripts, read length (~28-nt) and read distribution across the transcripts were reminiscent of sequencing data from ribosome profiling studies^31^. Second, the loss of 5’ tRNA halves in EDTA and ACD samples^6,13^ is consistent with loss of protection by the RNA-binding protein ZNF598 after polysome disassembly due to Mg2+ chelation. We have recently shown that ZNF598 binds tRNAs and translating ribosomes^32^, and the circulating tRNA halves correspond precisely to the region protected by the ZNF598. Chelation of Mg2+ by EDTA, traditionally used experimentally for that purpose^25,33^, and ACD in blood collection tubes will lead to disassembly of polysomes render the associated tRNAs vulnerable to nuclease digestion. The more widespread effect of RNP destabilization after Mg2+ chelation is furthermore evident by loss of RNA fragments from certain regions of U1 RNA, and overall fewer captured transcripts in EDTA and ACD samples though it ultimately remains unclear how much ex vivo effects of the different plasma additives on hematopoietic cells contribute to these differences^21–23^. Aside from the utility to study in vivo changes of exRNAs and to develop diagnostic applications, the discriminatory value of ex-mRNA compared to other RNA classes can also be utilized to assess such changes and biases related to blood collection and processing, which are well known in laboratory medicine and of which the effects of EDTA and ACD are the most prominent.

The finding that most reads originate from the coding sequence and not the UTRs is in contrast to a recent report by Skog *et al*.^15^ and likely due to different sRNA-seq protocols and analysis strategies. In fact, while Skog *et al*. and Danielson *et al*.^14^ used RNA end-treatment with RNA-seq they did not report enrichment of ex-mRNAs. In our study, strict mapping criteria were necessary to increase the signal-to-noise ratio for ex-mRNAs.

The adoption of exRNAs as clinical biomarkers will require quantitative and reasonably fast assays like qPCR. However, primer design for short fragments is challenging, and qPCR like other not-sequencing based assays does not easily allow to verify the amplified signal (i.e. read sequence). The diminutive amounts of RNA in body fluids increases the risk of introducing biases. For instance, up to 30% of reads in samples not end-treated with T4 PNK in this study mapped to the plasmid of Rnl2 ligase, which is used for adapter ligation during the sRNA-seq cDNA preparation. Omitting this plasmid reference from the mapping hierarchy during sequence read alignments resulted in a substantial amount of plasmid sequences aligning perfectly to other RNA classes, including mRNA transcripts, even using the most stringent mapping criteria. Similar considerations will have to be taken into account with different methods or further refinements, like e.g. using heparinase treatment to reduce possible interference of heparin with enzymatic reactions, or enzymatic rRNA removal.

In conclusion, total exRNA PNK-treatment followed by sRNA-seq allows for robust investigation of ex-mRNA changes for biomarker discovery and other studies. Future method refinements, such as depletion of rRNA and tRNA fragments, will further increase the potential of this approach.

## Acknowledgements

We thank the Rockefeller Genomics Resource Centre for performing the sequencing, and Tasos Gogakos from the Tuschl laboratory for scientific input and helpful discussions.

---

**Supplementary Fig. 1.**
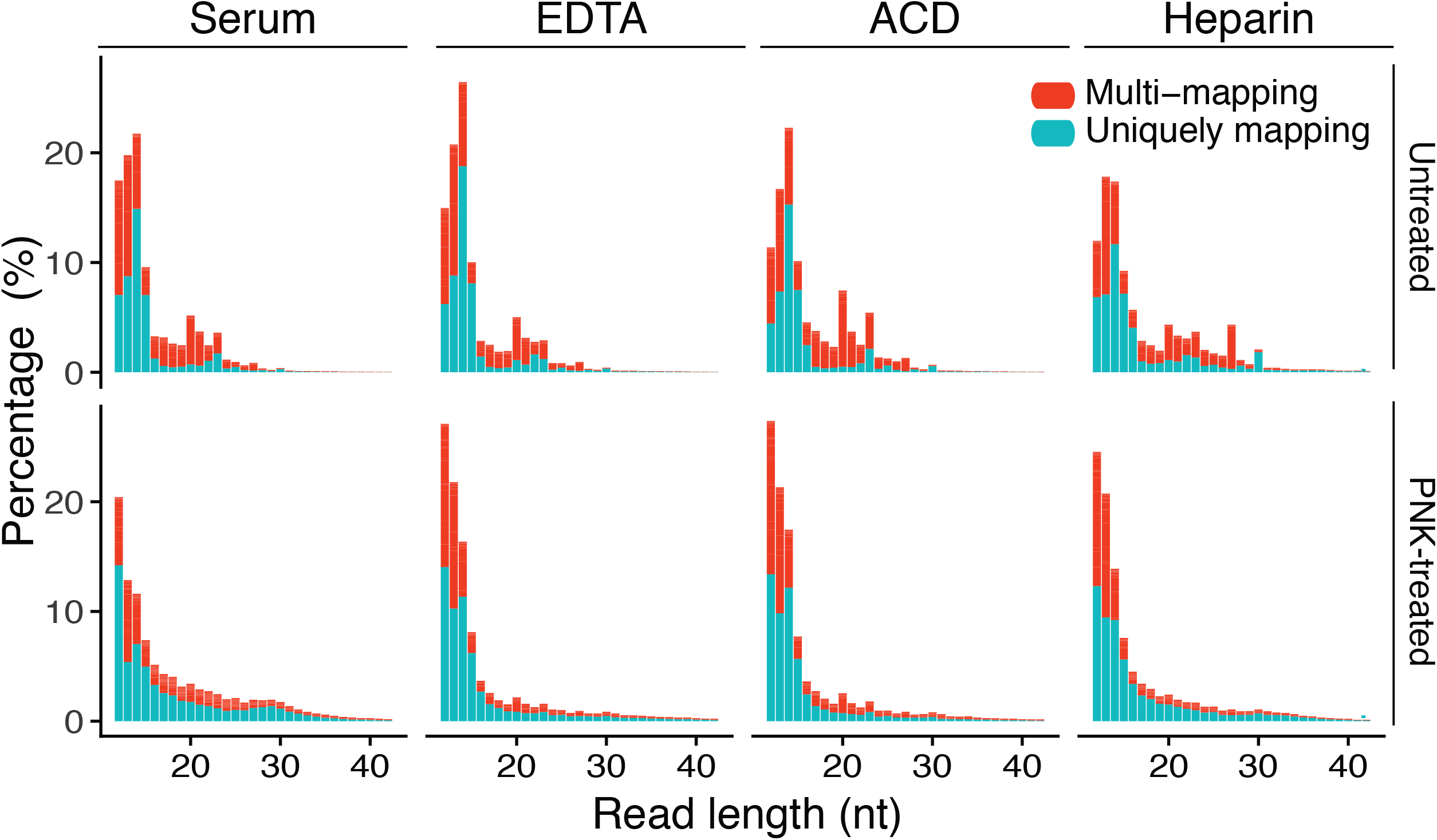

**Supplementary Fig. 2.**
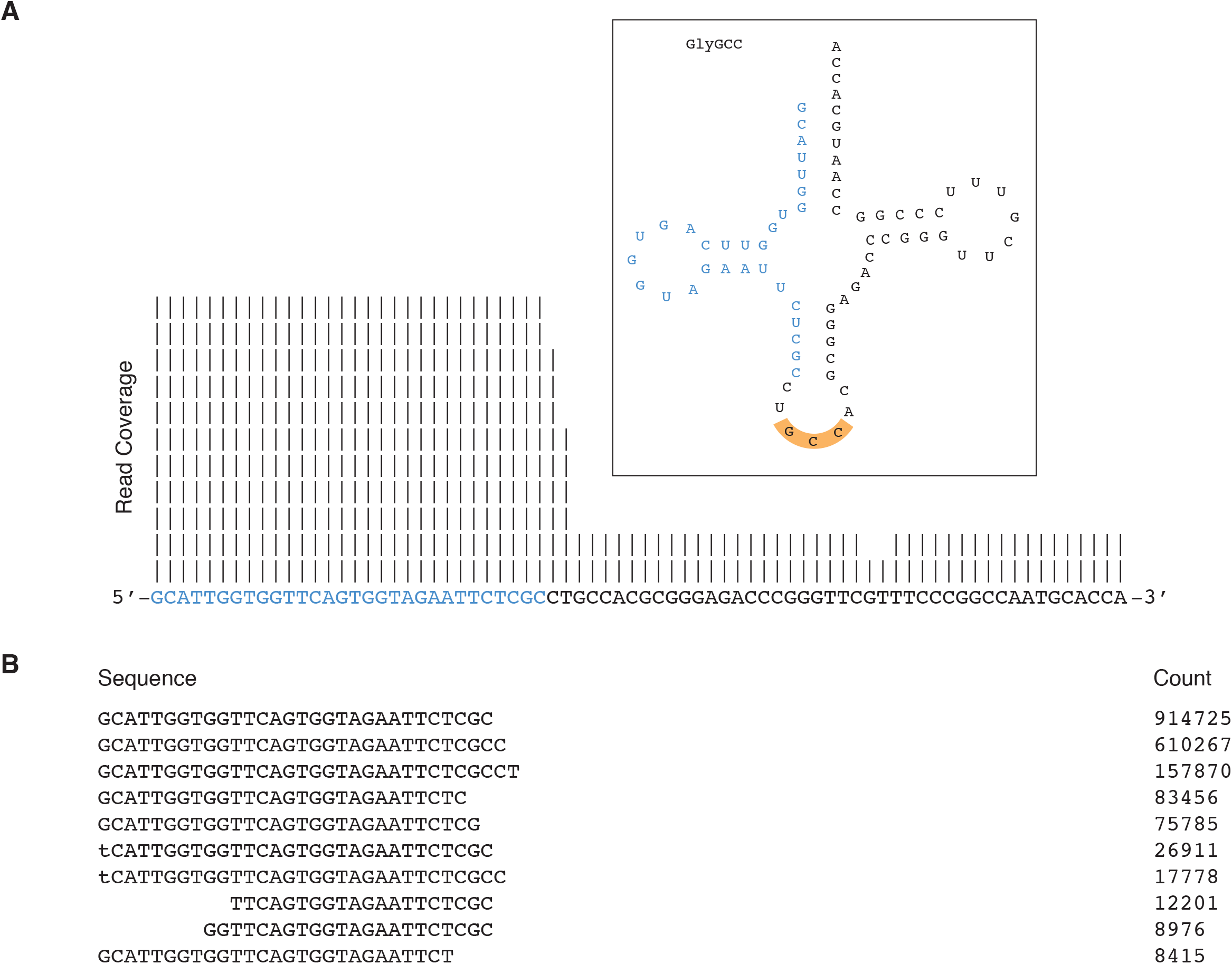

**Supplementary Fig. 3.**
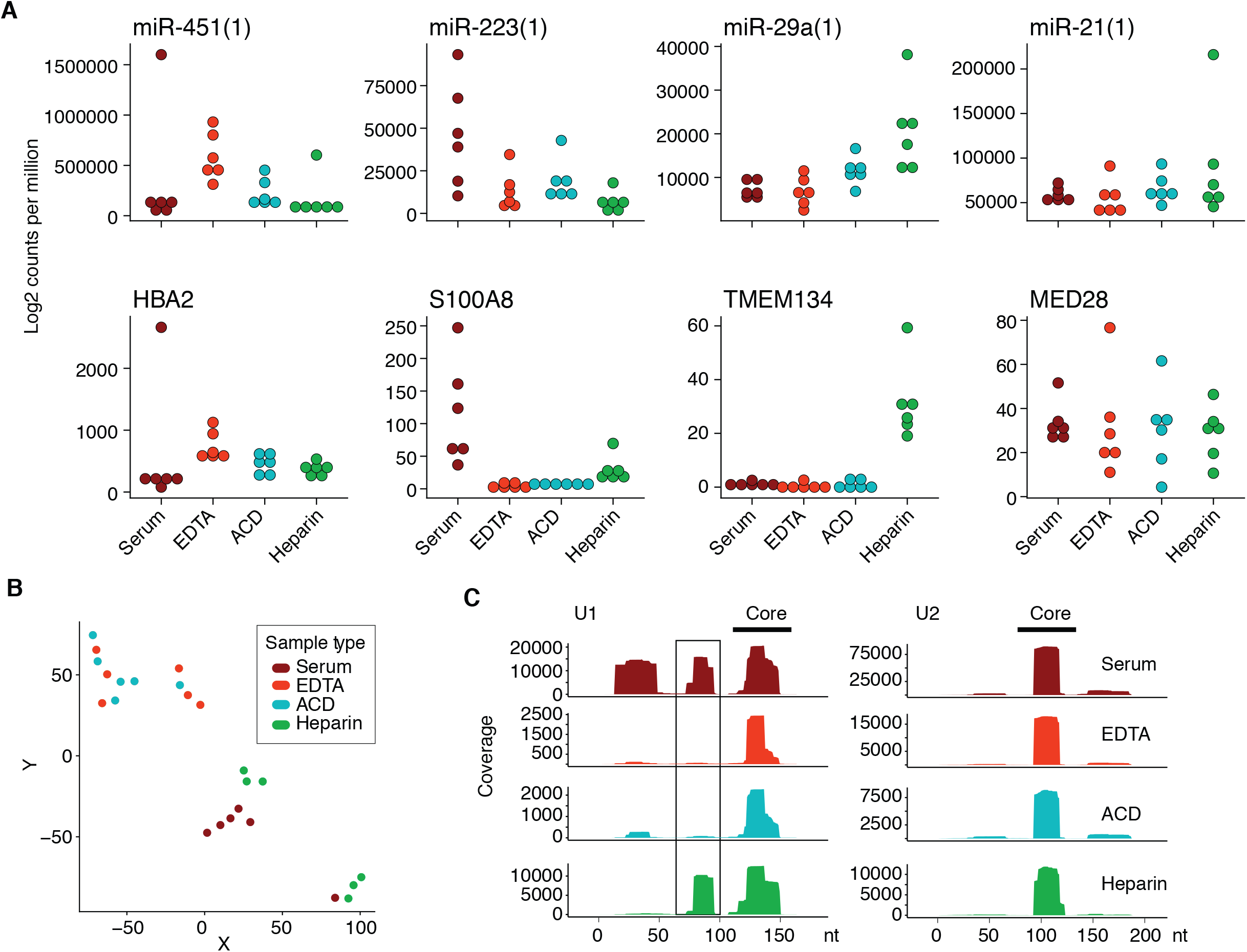

**Supplementary Fig. 4.**
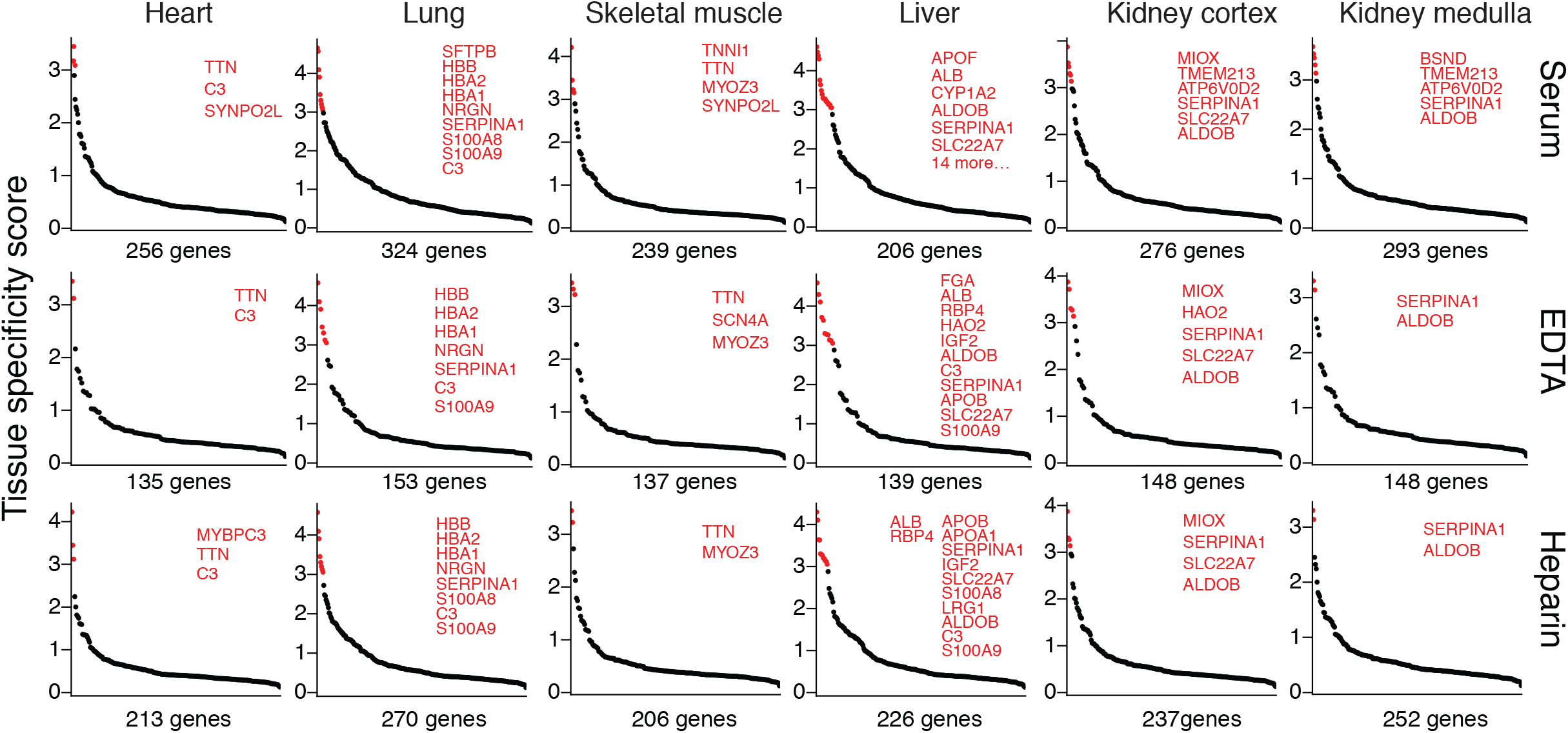

**Supplementary Fig. 5.**
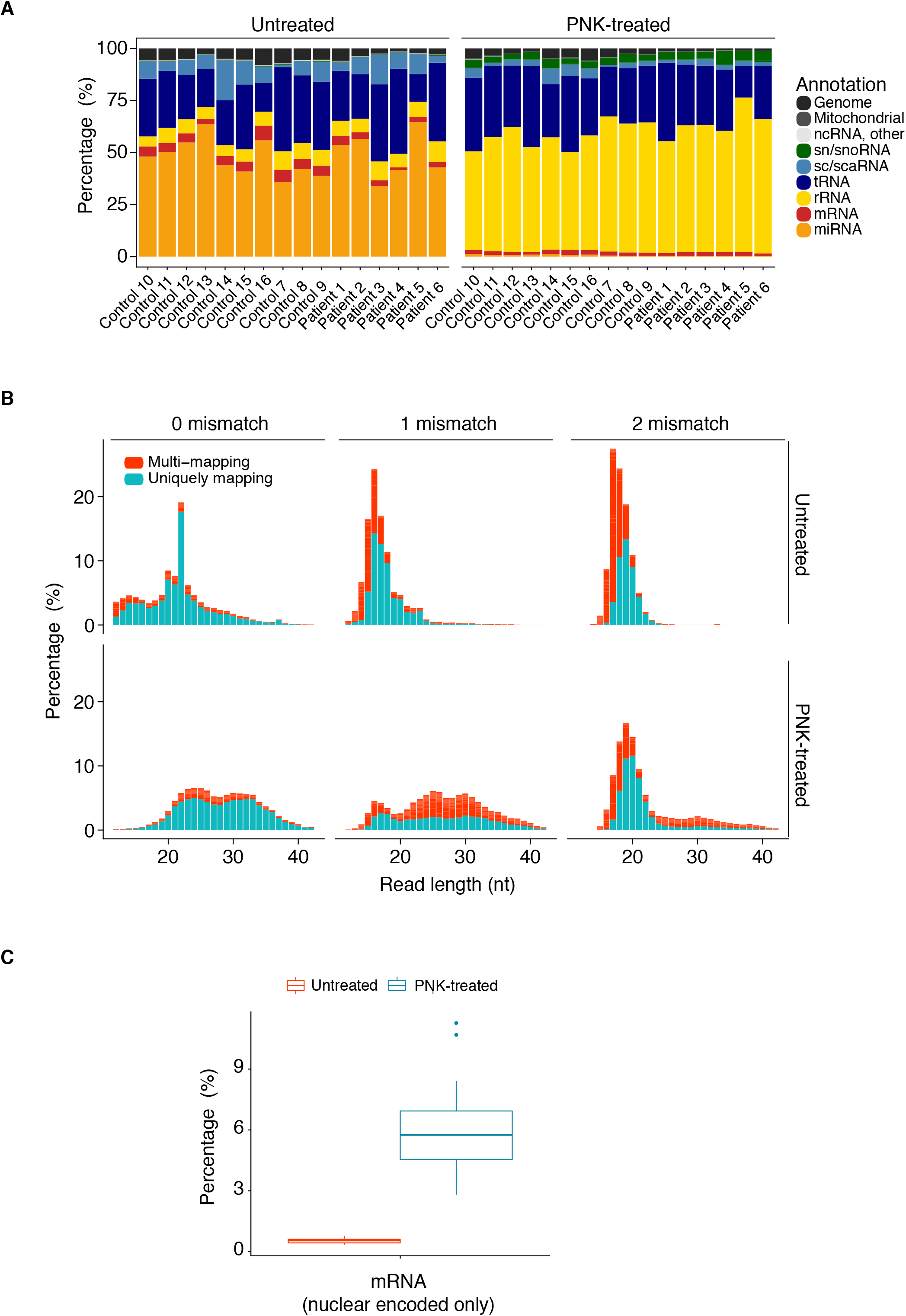

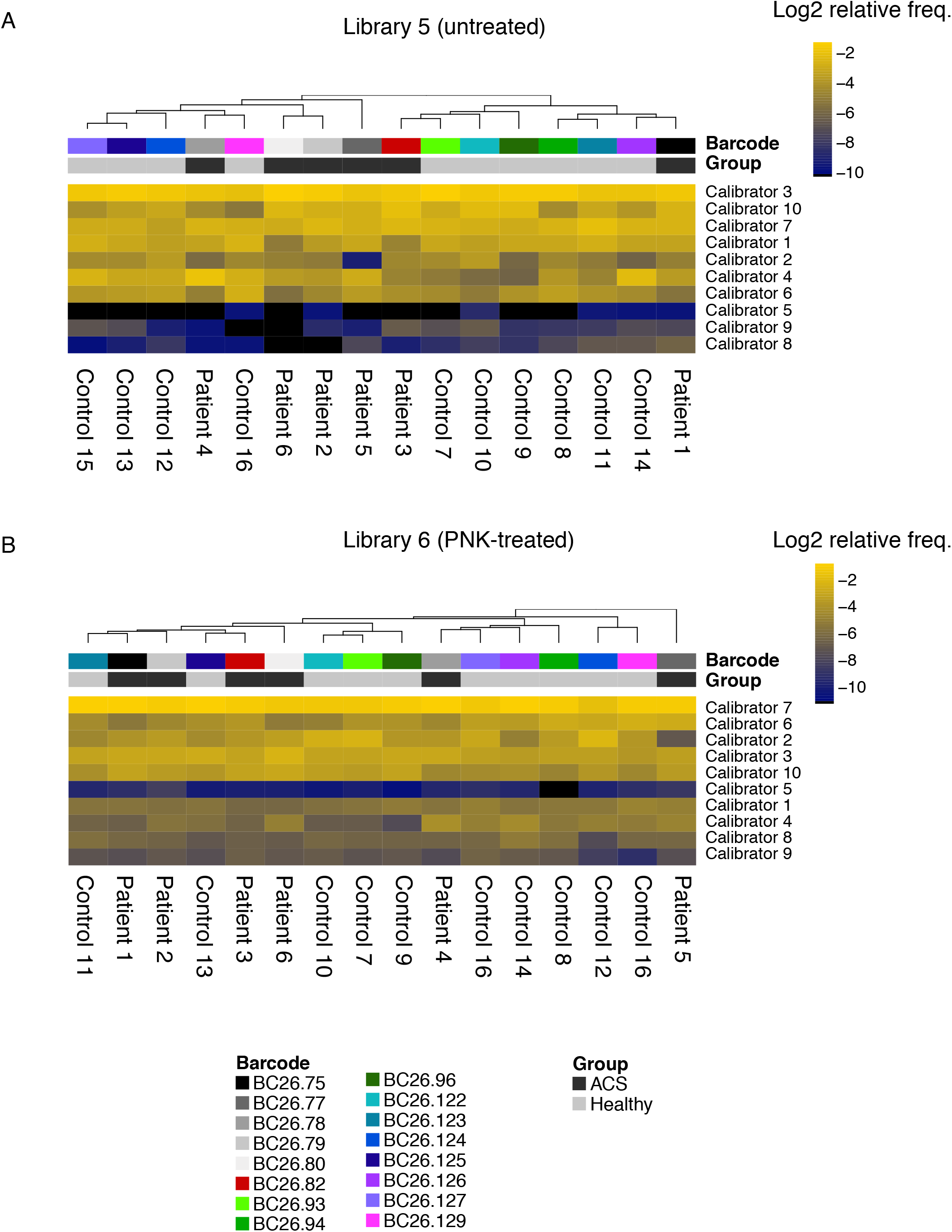

## References

1. Mandel, P. & Métais, P. Les acides nucléiques du plasma sanguin chez l’homme. C R Acad Sci III 142, 241–243 (1948).

2. Mitchell, P. S. et al. Circulating microRNAs as stable blood-based markers for cancer detection. Proc Natl Acad Sci U S A 105, 10513–10518 (2008).

3. Arroyo, J. D. et al. Argonaute2 complexes carry a population of circulating microRNAs independent of vesicles in human plasma. Proc Natl Acad Sci U S A 108, 5003–5008 (2011).

4. Turchinovich, A., Weiz, L., Langheinz, A. & Burwinkel, B. Characterization of extracellular circulating microRNA. Nucleic Acids Res 39, 7223–7233 (2011).

5. Elkayam, E. et al. The Structure of Human Argonaute-2 in Complex with miR-20a. Cell 150, 100–110 (2012).

6. Akat, K. M. et al. Comparative RNA-sequencing analysis of myocardial and circulating small RNAs in human heart failure and their utility as biomarkers. Proc Natl Acad Sci U S A 111, 11151–11156 (2014).

7. Corsten, M. F. et al. Circulating MicroRNA-208b and MicroRNA-499 reflect myocardial damage in cardiovascular disease. Circ Cardiovasc Genet 3, 499–506 (2010).

8. Max, K. E. A. et al. Human plasma and serum extracellular small RNA reference profiles and their clinical utility. Proc Natl Acad Sci U S A 115, E5334–E5343 (2018).

9. Landgraf, P. et al. A mammalian microRNA expression atlas based on small RNA library sequencing. Cell 129, 1401–1414 (2007).

10. Gerstberger, S., Hafner, M. & Tuschl, T. A census of human RNA-binding proteins. Nat Rev Genet 15, 829–845 (2014).

11. Giraldez, M. D. et al. Comprehensive multi-center assessment of small RNA-seq methods for quantitative miRNA profiling. Nat Biotechnol (2018).

12. Willeit, P. et al. Circulating MicroRNAs as Novel Biomarkers for Platelet Activation. Circ Res 112, 595–600 (2013).

13. Dhahbi, J. M. et al. 5’ tRNA halves are present as abundant complexes in serum, concentrated in blood cells, and modulated by aging and calorie restriction. BMC Genomics 14, 298 (2013).

14. Danielson, K. M., Rubio, R., Abderazzaq, F., Das, S. & Wang, Y. E. High Throughput Sequencing of Extracellular RNA from Human Plasma. PLoS One 12, e0164644 (2017).

15. Skog, J. et al. Glioblastoma microvesicles transport RNA and proteins that promote tumour growth and provide diagnostic biomarkers. Nat Cell Biol 10, 1470–1476 (2008).

16. Hafner, M. et al. Barcoded cDNA library preparation for small RNA profiling by next-generation sequencing. Methods 58, 164–170 (2012).

17. Farazi, T. A. et al. Bioinformatic analysis of barcoded cDNA libraries for small RNA profiling by next-generation sequencing. Methods 58, 171–187 (2012).

18. Dobin, A. et al. STAR: ultrafast universal RNA-seq aligner. Bioinformatics 29, 15–21 (2013).

19. Liao, Y., Smyth, G. K. & Shi, W. featureCounts: an efficient general purpose program for assigning sequence reads to genomic features. Bioinformatics 30, 923–930 (2014).

20. Wei, Z. et al. Coding and noncoding landscape of extracellular RNA released by human glioma stem cells. Nat Commun 8, 1145 (2017).

21. Brunialti, M. K., Kallás, E. G., Freudenberg, M., Galanos, C. & Salomao, R. Influence of EDTA and heparin on lipopolysaccharide binding and cell activation, evaluated at single-cell level in whole blood. Cytometry 50, 14–18 (2002).

22. Duvigneau, J. C. et al. Heparin and EDTA as anticoagulant differentially affect cytokine mRNA level of cultured porcine blood cells. J Immunol Methods 324, 38–47 (2007).

23. Shalekoff, S., Page-Shipp, L. & Tiemessen, C. T. Effects of anticoagulants and temperature on expression of activation markers CD11b and HLA-DR on human leukocytes. Clin Diagn Lab Immunol 5, 695–702 (1998).

24. Ukita, T., Terao, T. & Irie, M. Inhibition of pancreatic ribonuclease-I activity by heparin. J Biochem 52, 455–457 (1962).

25. Reveillaud, I., Lelay-Taha, M. N., Sri-Widada, J., Brunel, C. & Jeanteur, P. Mg2+ induces a sharp and reversible transition in U1 and U2 small nuclear ribonucleoprotein configurations. Mol Cell Biol 4, 1890–1899 (1984).

26. Zeller, T. et al. Assessment of microRNAs in patients with unstable angina pectoris. Eur Heart J 35, 2106–2114 (2014).

27. Karakas, M. et al. Circulating microRNAs strongly predict cardiovascular death in patients with coronary artery disease-results from the large AtheroGene study. Eur Heart J 38, 516–523 (2017).

28. Ward, J. A. et al. Circulating microRNA profiles in human patients with acetaminophen hepatotoxicity or ischemic hepatitis. Proc Natl Acad Sci U S A 111, 12169–12174 (2014).

29. Franck, G. et al. Flow Perturbation Mediates Neutrophil Recruitment and Potentiates Endothelial Injury via TLR2 in Mice: Implications for Superficial Erosion. Circ Res 121, 31–42 (2017).

30. Lee, T. D. et al. CAP37, a novel inflammatory mediator: its expression in endothelial cells and localization to atherosclerotic lesions. Am J Pathol 160, 841–848 (2002).

31. Ingolia, N. T., Ghaemmaghami, S., Newman, J. R. & Weissman, J. S. Genome-wide analysis in vivo of translation with nucleotide resolution using ribosome profiling. Science 324, 218–223 (2009).

32. Garzia, A. et al. The E3 ubiquitin ligase and RNA-binding protein ZNF598 orchestrates ribosome quality control of premature polyadenylated mRNAs. Nat Commun (2017).

33. Penman, S., Vesco, C. & Penman, M. Localization and kinetics of formation of nuclear heterodisperse RNA, cytoplasmic heterodisperse RNA and polyribosome-associated messenger RNA in HeLa cells. J Mol Biol 34, 49–60 (1968).

