## Supplementary Data 9 for "Detection of circulating extracellular mRNAs by modified small RNA-sequencing analysis"

|  | ACS | Controls | P value |
| --- | --- | --- | --- |
| n | 6 | 10 |  |
| Age - mean (sd) | 62 (8.93) | 54 (5.64) | 0.056 |
| Hb - mean (sd) | 13.8 (2.02) | 14.7 (0.83) | 0.24 |
| WBC - mean (sd) | 9.2 (2.22) | 5.9 (1.00) | 0.001 |
| PLT - mean (sd) | 256 (58) | 237 (65) | 0.575 |
