## Supplementary Information for "Detection of circulating extracellular mRNAs by modified small RNA-sequencing analysis"

### Detailed Methods

#### Sample collection

Peripheral vein blood was collected from healthy volunteers at The Rockefeller University Hospital (IRB number KAK-0839), and from patients with acute coronary syndrome at the First Department of Medicine and the Department of Emergency Medicine at the University Hospital Mannheim (IRB: protocols KAK-0750 and 2009-300N-MA). Human tissue samples for bulk mRNA-seq of myocardium and kidney were obtained from the National Disease Research Interchange (Philadelphia), and other samples were obtained from biopsies or discarded surgical waste (Rockefeller University IRB number TTU-0707). Sample procurement was approved by the institutional review boards of all participating institutions. All participants gave written informed consent.

#### Serum and plasma sample processing and platelet-depletion

Serum samples were allowed to coagulate before further processing; plasma samples were processed within 30 min of collection by centrifugation at 2500 x g for 15 min in the blood collection tube, followed by another centrifugation of the supernatant at the same conditions. The resulting supernatant was aliquoted into 500 µl aliquots avoiding the residual pellet, flash-frozen in liquid nitrogen and stored at -80 °C until use. Samples were thawed only once.

The first author collected or supervised the blood collection from an antecubital vein at both sides, and processed all samples.

#### RNA isolation

##### Isolation of total cell-free, extracellular RNA (exRNA)

Total exRNA was isolated from 450 µl biofluid using a customized RNA isolation protocol that we developed and recently published to quantitatively inhibit nuclease activity ^1^.

##### Isolation of cell and tissue RNAs

Cellular or tissue total RNA was extracted using TRIzol with an additional phenol/chloroform extraction step and concentrated by alcohol precipitation or purified using the miRNeasy kit (Qiagen) according to the manufacturer’s instructions.

#### RNA quantification and quality control

Total exRNA from selected samples was quantified using the Qubit RNA HS Assay Kit (ThermoFisher, cat# Q32852) on a Qubit 1.0 Fluorometer (ThermoFisher) as described ^1^.

Total RNA from cells and tissues was quantified on a NanoDrop UV spectrophotometer, and the RNA integrity was determined on an Agilent Bioanalyzer 2100 with Agilent RNA 6000 Pico Chips.

#### PNK treatment of total exRNA

The untreated total exRNA after the initial isolation (above) was eluted with 30 µl RNase-free water, resulting in about 28 µl total. To half of it, 14 µl, we added 6 µl of a master mix corresponding to the equivalent of 2 µl 10x T4 PNK buffer, 2 µl 10 mM ATP, 1 µl RNase-free water, and 1 µl T4 polynucleotide kinase (PNK, NEB, cat# M0201S) for a final reaction volume of 20 µl in a 1.5 ml microcentrifuge tube. The reaction was incubated for 30 min at 37 °C followed by the addition of 40 µl buffer VB2G (for details see Max et al.^1^) and reapplied to the same silica-column used for the initial purification. The column was washed twice with 900 µl buffer EWL (for details see Max *et al.*^1^), once with 900 µl 100% ethanol, and once with 500 µl 80% ethanol. The column was then placed in a collection tube and spun at 13000 rpm and room temperature for 5 min to dry the silica matrix. The silica column was then transferred to a new 1.5 ml tube, 15 µl RNase-free water was applied to the column, incubated for 1 min and finally centrifuged at 13000 rpm and room temperature for 1 min to elute the PNK-treated RNA. This yielded ~13 µl and 8.5 µl was used for sRNA-seq cDNA library preparation.

#### Small RNA-sequencing

We performed barcoded small RNA-sequencing (sRNA-seq) as published previously ^2^ but with additional barcoded 3’-adapters to allow multiplexing of up to 24 samples; the detailed number of samples per library and the number used in this study is indicated in Supplementary Data I. The cDNA library preparation started with a 3’ ligation of a pre-adenylated DNA oligonucleotide for each individual sample. The 3’-adapter master mix contained a cocktail of 10 equimolar 22-nt synthetic RNAs (Supplementary Data XIV, cocktail 2). The samples were pooled after the 3’ adapter ligation, size selected for small RNAs but the upper limit was chosen to be 45-nt instead of 24-nt used for classic miRNA-seq to allow longer mRNA fragments to be included; that is the size selection was 19- to 45-nt. RNA ligated to the 3’ adapter was gel purified, followed by 5’ ligation of an RNA oligonucleotide and another size selection and gel purification. The cDNA library preparation was completed by second strand synthesis using SuperScript III, and PCR amplification. The resulting amplicon was quality controlled on an Agilent TapeStation with a High Sensitivity D1000 ScreenTape and used for cluster generation and sequencing 50-bp single-read on a HiSeq 2500 sequencer in rapid run mode.

#### mRNA-sequencing

mRNAseq libraries were prepared by the Genomics Core Facility of The Rockefeller University using the Illumina TruSeq Stranded mRNA LT protocol according to the manufacturer’s protocol but using NEB’s Protoscript II reverse transcriptase for the first-strand cDNA synthesis. RNA input was 400 ng total RNA. Individual RNAseq libraries were quality controlled on an Agilent TapeStation with a High Sensitivity D1000 ScreenTape. The libraries were sequenced on an Illumina NextSeq 500 sequencer 75-bp paired-end in high-output mode in the Genomics Core Facility of The Rockefeller University.

#### Bioinformatics analysis

##### FASTQ data file generation

Raw image data were converted to FASTQ files by the Genomics Core Facility of The Rockefeller University using Illumina’s bcl2fastq conversion software version 2.18.0.12 (NextSeq 500) or version 1.8.4 (HiSeq 2500).

##### sRNA-seq read annotation

Short reads were annotated as described^3,4^. FASTQ files were collapsed and demultiplexed followed by alignment with the BWA aligner^5^ to a combined transcriptome reference and the unmapped reads to a combined genome reference. Mapping parameters were set to allow for a maximum of 2 mismatches and reporting up to 999 alignments. The transcriptome reference consisted of spike-in sequences for oligonucleotides used during the cDNA library preparation (Supplementary Data XIV), processed mRNAs (Ensembl release 75), rRNAs, miRNAs, tRNAs, small nuclear (sn) and small nucleolar (sno) RNAs, small cytoplasmic (sc) and small Cajal-bodies (sca) RNAs, and other minor non-coding RNA categories. The genome reference consisted of the human genome reference build 38 (primary assembly) and the E. coli K-12 genome.

After the initial read alignment the final annotation category was assigned in a hierarchical manner^3,4^ to give RNA transcript categories with higher abundance or higher likelihood of being the true category precedence over less abundant transcripts. The highest priority had synthetic RNA transcripts, introduced intentionally like spiked-in calibrators or unavoidable contaminants in recombinant enzyme preparations. The applied hierarchy was (descending priority):

CalibratorIllumina

CalibratorTuschlSet1

CalibratorTuschlSet2

CalibratorTuschlSet3

CalibratorTuschlSet4

Adapter

SizeMarker

Plasmid

EColiK12

Diatom

rRNA

MTrRNA

tRNA

MTtRNA

snRNA

snoRNA

MTmRNA

scRNA

scaRNA

miRNA

mRNA

circRNA

lincRNA

rRNAPrecursor

tRNAPrecursor

snRNAPrecursor

snoRNAPrecursor

scRNAPrecursor

scaRNAPrecursor

piRNA

miRNALowConfidence

snoRNALowConfidence

MTGenome

Genome

Viral

With that approach a read mapping to an rRNA transcript with 2 mismatches and an mRNA transcript without mismatch will be finally annotated as rRNA, the category higher in the hierarchy. miRNA nomenclature

To reduce redundancy, individual miRNAs with identical mature, i.e. ~22-nt, sequences were reported with the number of members in parenthesis, e.g. miR-122(1) for one member miRNAs, miR-194(2) for two member miRNA, and so forth.

##### mRNA read annotation

The mRNAseq reads were aligned to the human genome build 38 using the STAR aligner^6^ (version 2.0.4j) allowing for two mismatches. Expression values (count matrices) were generated using the featureCounts application from the Rsubread^7^ software package (version 1.5.1) and gene definitions from Ensembl release 82.

##### Differential analysis

Differential analysis of miRNA and mRNA differences was done with the Bioconductor package *edgeR (*version edgeR_3.17.5*)* in the R statistical language ^8^. We used edgeR’s quasi-likelihood (QL) framework ^9,10^ to fit a simple generalized linear model comparing the conditions of interest. The QL dispersion distribution was estimated robustly, other parameters were kept at their default setting. miRNAs or mRNAs with fewer than 1 and 0.5 counts per million, respectively, in the number of samples comprising the smallest group for each comparison were excluded from the differential analysis and all downstream analyses. The mroast function^11^was used for gene set analysis. Other statistical tests are indicated in text and figures where appropriate. An FDR of < 10% was used as cut-off.

##### Metagene analysis

To determine if the mRNA read distribution was biased towards certain transcript regions, i.e. 5’ UTR, CDS and 3’ UTR, we compared the experimental data with 100 random simulations. We only considered mRNA transcripts with UTR’s 15-nt or longer, and RNA-seq reads 15-nt or longer with only 2 mapping locations to the transcriptome. The reads were taken from each subsample (i.e. 6) per sample type. The selected reads were mapped using the bowtie aligner (version 1) to the subset transcriptome. The mapped reads for each region per distinct transcript were tallied and normalized by region length and averaged across all transcripts. Finally, the transcript regions were scaled to 100%. For the simulation, the transcript region mapping positions of the previously mapped reads were randomly shuffled and counted as above. This simulation was repeated 100 times and the results were averaged.

##### Tissue specificity score

Entropy as a measure of tissue-specific expression (tissue-specificity score) was calculated as described^12^. The selected samples/their identifiers are given in Supplementary Data VII.

### Supplementary Figure Legends

**Supplementary Fig. 1. Read length distribution of uniquely mapping and multi-mapping mRNAs reads across different blood sample types.** Included were only reads without mismatch to the reference transcriptome.

**Supplementary Fig. 2. Read coverage of ex-tRNAs.** (A) Read coverage of the Gly-tRNA isoacceptor from the untreated heparin sample of donor “Control 1” (sample “Control1_heparin_untreated” in Supplementary Data 1) together with the tRNA secondary structure (box). The anticodon is highlighted in yellow and bases protected by the RNA-binding protein ZNF598 based on PAR-CLIP^13^ are colored blue. The pileup was scaled to 11 bins using the following formula for each base: (((count/maximum count) ** 1/4) + 0.5)/0.1. (B) Showing the 10 most frequently sequenced sequences with corresponding read count.

**Supplementary Fig. 3. Differences in the extracellular miRNA and mRNA profile and read coverage by blood sample type.** (A) Differences in abundance of cell-free, circulating miRNAs (top row) and mRNAs (bottom row); see Supplementary Data XIV for hypothesis tests. (B) Read coverage for the small nuclear RNAs (snRNAs) U1 (left), and U2 (right) in the different sample types. Rectangle in the left sub-figure indicates coverage loss of U1 region in EDTA and ACD plasma. Bar (core) on the left (U1) and right (U2) indicates core region resistant to nuclease digestion. (C) t-SNE plot based on the exRNA profiles of all RNA categories of the four different sample types from six healthy donors (perplexity 4).

**Supplementary Fig. 4.** **Capture of top expressed transcripts from selected solid tissues in different sample types.** The 1,000 most abundant nuclear mRNA transcripts from the selected cell types and tissues from bulk RNA-seq that collected 5 unique or 10 total ex-mRNA reads in at least 3 of the 6 donors per sample type were considered captured. The x out of 1,000 captured transcripts (x axis) were ordered in descending order by the tissue specificity score (TSS, y axis). Transcripts with a TSS greater than 3 were highlighted in red and listed space permitting. Note that bulk sequencing data of heterogenous and well perfused tissues like lung will contain hematopoietic or erythropoietic/hematopoietic-specific transcripts within the top 1,000 genes of the sample. It should also be noted that most reads for e.g. the MYBPC3 (left lower panel) and all for MIOX (kidney panels) were shorter than 17-nt with a high likelihood of misassignments.

**Supplementary Fig. 5**. Treatment of total extracellular RNA with T4 polynucleotide kinase (T4 PNK) followed by small RNA-sequencing (sRNA-seq) in a pilot cohort of chest pain patients (ACS) and matched controls. This figure is analogous to Figure 1B, C and Supplementary Fig. I showing basic read and annotation characteristics (A, B) as well as the effect of PNK end-treatment on mRNA capture (C) for libraries 5 and 6. (A) Differences in read annotation for endogenous RNA classes for untreated RNA and PNK-treated RNA using initial annotation settings (up to 2 mismatches, unlimited multi-mapping). (A) Read length distribution for reads annotated as mRNAs (nuclear and mitochondrial) with 0, 1, and 2 mismatches to the reference transcriptome; multi-mapping reads and reads mapping to only one transcript (uniquely mapping) are shown in different colors as indicated. Note that for the final mRNA downstream analysis only mRNA reads without mismatch (i.e. 0 mismatch), maximum 2 mapping locations, and 15-nt or longer were considered. (C) Differences in nuclear-encoded mRNA capture between untreated and PNK-treated RNA using strict (final) annotation criteria (no mismatch and up to two mapping locations).

### Supplementary Data

**Supplementary Data 1.** Master table in Excel spreadsheet format containing each sample with sRNA-seq library information and read annotation as per initial annotation settings (up to two mismatches, multi-mapping allowed) shown as read counts and percentages.

**Supplementary Data 2**. Effect of mismatches and multi-mapping on the percentage of reads annotated as mRNAs.

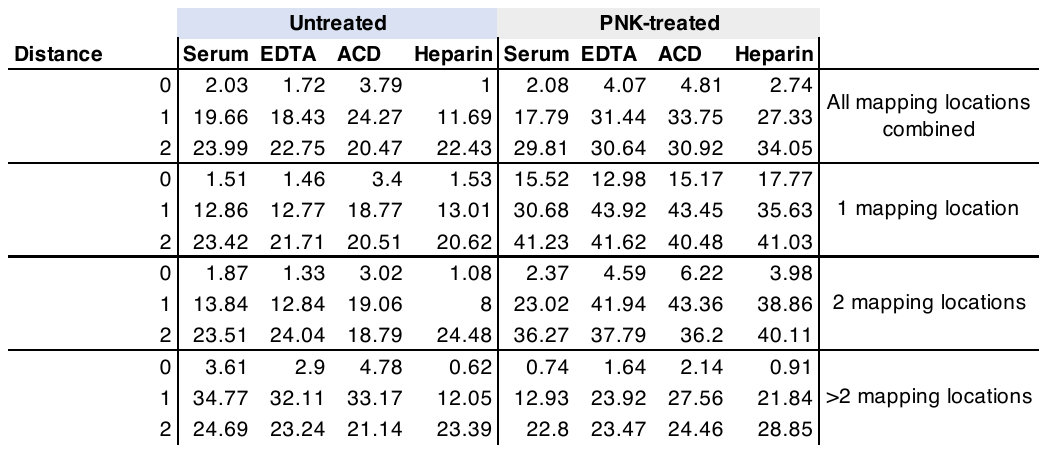

**Supplementary Data 3.** Excel spreadsheet containing the results of the differential analysis of circulating miRNAs comparing the 4 different sample types used in libraries 1 to 4. Shown are raw and Benjamini-Hochberg corrected (FDR) P values for the overall/ANOVA-like analysis and Benjamini-Hochberg corrected (FDR) P values of the pairwise comparisons.

**Supplementary Data 4**. Excel spreadsheet containing the results of the differential analysis of circulating mRNAs comparing the 4 different sample types used in libraries 1 to 4. Only mRNA fragments without mismatch, 15-nt or longer, and at maximum mapping to two transcripts of the transcriptome reference were allowed for this analysis. Shown are raw and Benjamini-Hochberg corrected (FDR) P values for the overall/ANOVA-like analysis (any difference among the samples?) and Benjamini-Hochberg corrected (FDR) P values of the pairwise comparisons.

**Supplementary Data 5**. Excel spreadsheet containing the results of the gene set analysis of mRNAs comparing the 4 different sample types used in libraries 1 to 4. C2 gene sets (from the MSigDB Collections of the Broad Institute) containing the terms “ribosome”, “translation”, or “inflammation” were used as input. A self-contained test as implemented in the mroast function of the Bioconductor package edgeR was used. The mRNA input count matrix and the design model were identical to the differential analysis (Dataset 3).

**Supplementary Data 6.** Excel spreadsheet containing the tissue and cell RNA-seq samples used to calculate tissue-specific expression (entropy/tissue-specificity score). Expression values are given in transcripts per million (TPM).

**Supplementary Data 7**. Excel spreadsheet containing the 144 mRNA genes (first column) with a tissue-specificity score > 3 confidently detected in circulation in either serum or the plasma samples EDTA, ACD, or heparin. The other columns show the corresponding expression values in TPM (transcripts per million) of these genes in the selected tissues.

**Supplementary Data 8.** Percentage the top 200, 500, 1,000 expressed transcripts in the shown cell and tissues captured confidently in circulation based on the sample type/anticoagulant.

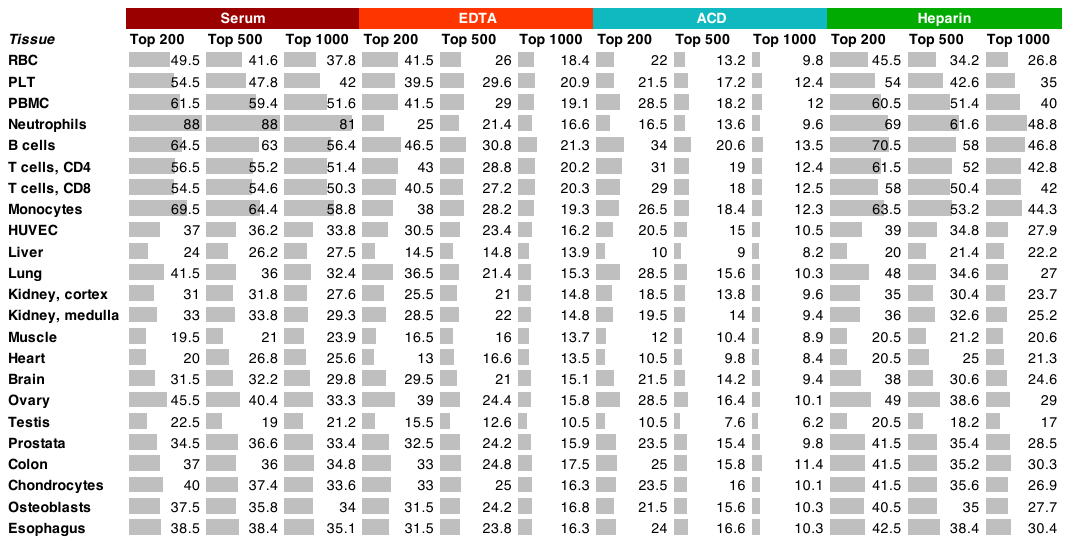

**Supplementary Data 9.** **ACS and control group demographics.** Basic demographics and blood parameters of 6 patients with acute coronary syndrome (ACS) and 10 matched controls used in libraries 5 (RNA untreated) and 6 (RNA PNK-treated). All 16 donors were male. P values calculated using the Wilcoxon rank sum test.

|  | ACS | Controls | P value |
| --- | --- | --- | --- |
| n | 6 | 10 |  |
| Age - mean (sd) | 62 (8.93) | 54 (5.64) | 0.056 |
| Hb - mean (sd) | 13.8 (2.02) | 14.7 (0.83) | 0.24 |
| WBC - mean (sd) | 9.2 (2.22) | 5.9 (1.00) | 0.001 |
| PLT - mean (sd) | 256 (58) | 237 (65) | 0.575 |

**Supplementary Data 10.** Excel spreadsheet containing the individual level clinical chemistry and laboratory data for the six ACS patients and the 10 matched controls whose RNA was processed in sRNA-seq libraries 5 (untreated) and 6 (PNK-treated).

**Supplementary Data 11**. Excel spreadsheet containing the results of the differential analysis of circulating miRNAs (library 5 only) comparing the 6 patients with troponin I-positive acute coronary syndrome (ACS) compared to age-matched controls.

**Supplementary Data 12**. Excel spreadsheet containing the results of the differential analysis of circulating mRNAs (library 6 only) comparing the 6 patients with troponin I-positive acute coronary syndrome (ACS) compared to age-matched controls. Only mRNA fragments without mismatch, 15-nt or longer, and at maximum mapping to two transcripts of the transcriptome reference were allowed for this analysis. Shown are raw and Benjamini-Hochberg corrected (FDR) P values for the overall/ANOVA-like analysis (any difference among the samples?) and Benjamini-Hochberg corrected (FDR) P values of the pairwise comparisons.

**Supplementary Data 13.** Sequences of the synthetic RNA cocktails used as spike-ins for the RNA isolation (cocktail 1), and the cDNA library generation (cocktail 2).

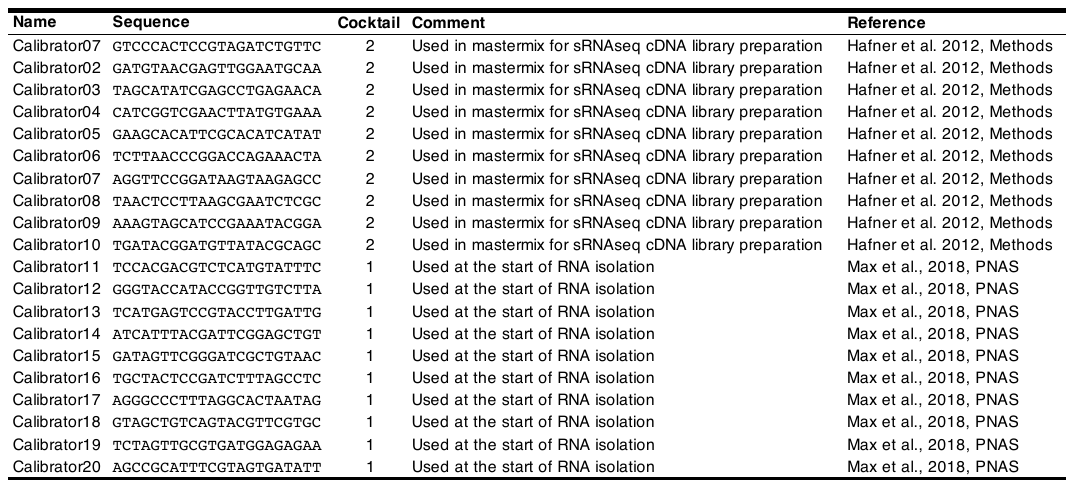

**Supplementary Data 14.** Benjamini-Hochberg corrected P values for pairwise comparisons as shown in Supplementary Fig. IIA.

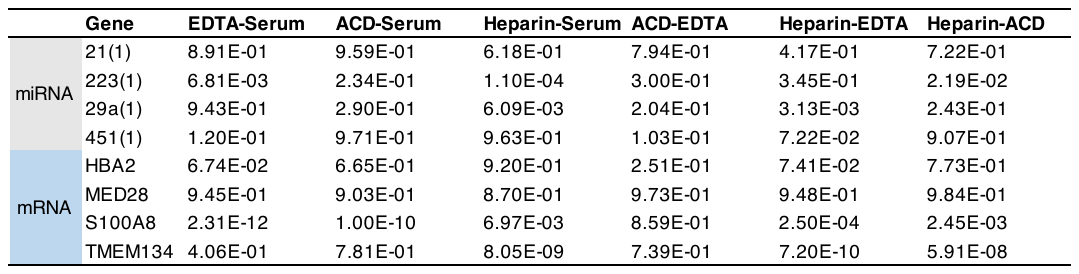

### Supplementary References

1.	Max, K. E. A. et al. Human plasma and serum extracellular small RNA reference profiles and their clinical utility. *Proc Natl Acad Sci U S A* **115**, E5334-E5343 (2018).

2.	Hafner, M. et al. Barcoded cDNA library preparation for small RNA profiling by next-generation sequencing. *Methods* **58**, 164-170 (2012).

3.	Brown, M., Suryawanshi, H., Hafner, M., Farazi, T. A. & Tuschl, T. Mammalian miRNA curation through next-generation sequencing. *Front Genet* **4**, 145 (2013).

4.	Farazi, T. A. et al. Bioinformatic analysis of barcoded cDNA libraries for small RNA profiling by next-generation sequencing. *Methods* **58**, 171-187 (2012).

5.	Li, H. & Durbin, R. Fast and accurate short read alignment with Burrows-Wheeler transform. *Bioinformatics* **25**, 1754-1760 (2009).

6.	Dobin, A. et al. STAR: ultrafast universal RNA-seq aligner. *Bioinformatics* **29**, 15-21 (2013).

7.	Liao, Y., Smyth, G. K. & Shi, W. featureCounts: an efficient general purpose program for assigning sequence reads to genomic features. *Bioinformatics* **30**, 923-930 (2014).

8.	R, D. C. T. R: A Language and Environment for Statistical Computing. (2014).

9.	Lun, A. T., Chen, Y. & Smyth, G. K. It’s DE-licious: A Recipe for Differential Expression Analyses of RNA-seq Experiments Using Quasi-Likelihood Methods in edgeR. *Methods Mol Biol* **1418**, 391-416 (2016).

10.	Robinson, M. D., McCarthy, D. J. & Smyth, G. K. edgeR: a Bioconductor package for differential expression analysis of digital gene expression data. *Bioinformatics* **26**, 139-140 (2010).

11.	Wu, D. & Smyth, G. K. Camera: a competitive gene set test accounting for inter-gene correlation. *Nucleic Acids Res* **40**, e133 (2012).

12.	Gerstberger, S., Hafner, M. & Tuschl, T. A census of human RNA-binding proteins. *Nat Rev Genet* **15**, 829-845 (2014).

13.	Garzia, A. et al. The E3 ubiquitin ligase and RNA-binding protein ZNF598 orchestrates ribosome quality control of premature polyadenylated mRNAs. *Nat Commun* (2017).
